# Single nuclei multiomic analyses identify human cardiac lymphatic endothelial cells associated with coronary arteries in the epicardium

**DOI:** 10.1101/2023.03.29.534454

**Authors:** Stanislao Igor Travisano, Michael R M Harrison, Matthew E. Thornton, Brendan H. Grubbs, Thomas Quertermous, Ching-Ling Lien

## Abstract

**Summary:** Cardiac lymphatic vessels play important roles in fluid homeostasis, inflammation, disease, and regeneration of the heart. The developing cardiac lymphatics in human fetal hearts are closely associated with coronary arteries, similar to those in zebrafish hearts. We identified a population of cardiac lymphatic endothelial cells that reside in the epicardium. Single nuclei multiomic analysis of the human fetal heart revealed the plasticity and heterogeneity of the cardiac endothelium. Furthermore, we found that VEGFC is highly expressed in arterial endothelial cells, providing a molecular basis for the arterial association of cardiac lymphatic development. Using a cell-type-specific integrative analysis, we identified a novel population of cardiac lymphatic endothelial cells marked by the PROX1, the lymphangiocrine RELN, and enriched in binding motifs of ETV transcriptional factors. We report the first in vivo molecular characterization of human cardiac lymphatics and provide a valuable resource to understand fetal heart development.

## Introduction

The first blood vessels form *de novo* by a vasculogenic process of endothelial precursors known as angioblasts. The newly assembled vascular bed grows and expands into a mature network through a process known as angiogenesis^1^. Blood and lymphatic vessel networks are tightly associated, share a similar endothelial organization but they are distinct at a molecular and structural level and function differently^2^. The lymphatic vessels collect and remove the interstitial fluid from the tissues and transport it to thoracic duct, returning it to the venous circulation^2^. Defects in lymphatic development or function have been associated to the progression of several immune and inflammatory diseases, as well as tumor metastasis^3^.

Dr. Florence Sabin suggested that endothelial cells (ECs) of embryonic veins sprout out as the lymphatic endothelial cells (LECs) and form the lymph sac^4^. However, the contribution of mesenchymal/hemogenic precursor cells (termed lymphangioblasts) to the development of the lymph sac independently from the veins has also been demonstrated by Huntington^5,6^. The idea of mesenchymal cells growing centripetally before creating an anastomosis with the venous system was further explored by Kampmeier^7,8^. His work on other mammals reinforced the idea that the lymphatic system develops from a mesenchymal cell population characterized by the presence of several vacuoles^9^. The centripetal and the centrifugal theories have been debated for over a century^10,11^. Although many works reported the venous origin of the lymphatic progenitors, only very recently has the discovery of new cell lineages contributing to organ-specific lymphatics changed the understanding of LEC origin^12^. New lines of evidence for non-venous sources were reported within the dermal^13^, intestine^14^, mesentery^15^, paraxial mesoderm^16^, cardiopharyngeal mesoderm^17^, and lastly, the cardiac second heart field (SHF)-derived lineage^18^.

The adult human cardiac lymphatics are made of two different plexuses within the subepicardium and the subendocardium as demonstrated by tissue clearing^19^ and this is also observed in most other mammals. However, the mechanism of how these two plexuses form is still unknown^19^. Anatomical study on the dog has shown that an extensive plexus of lymphatic vessels is located within the epicardium of the beating heart^20^. Moreover, these subepicardial cardiac lymphatics follow the branches of the coronary artery (CA)^21^. Interestingly in contrast to reports in mice^22^ where cardiac lymphatics are mainly along the coronary veins, we observed that cardiac lymphatics on the surface of zebrafish hearts are mostly associated with CA which serve as a scaffold for the elongation of the lymphatic vessels^23,24^. Furthermore, it is known that lymphatic vessels in different regions of skin might develop by different mechanisms with or without obvious alignment with blood vessels^13^. The anatomical differences in the development of the lymphatic vessels suggest potential organ or species specific sources of the lymphatic progenitors^25^. It remains unknown whether that might account for the different associations of the lymphatic vessels with arteries or veins.

The epicardium is a mesothelial monolayer of cells that cover the entire heart in all vertebrates and contributes to cells in the coronary vasculature and connective tissue of the heart ^26^. In human, the epicardium develops between Carnegie Stage 12-16^27,28^, while the embryonic coronary vasculature of the human heart begins to form as a group of epicardial blood islands, endothelium-lined cysts filled with nucleated erythrocytes^29^. In the epicardium, angioblasts organize dorsally in tubular structures and migrate into the myocardium to form similar structures, via vasculogenesis^30^. These cells, as visualized with electron microscopy, can be seen in three configurations: blood island-like structures with erythrocytes, forming tubes by joining with other angioblasts, and in the formation of a large vacuole, which displaces their cytoplasm^30^. In mice, two independent studies identified resident macrophage population that reside in the subepicardial compartment and regulates the emergence of new lymphatics^31^, and a SHF derived, sub-mesothelial cell population that resides at the base of the pulmonary artery and has the ability to differentiate into cardiac lymphatic^32^. The anatomy and presence of different progenitor populations of human cardiac lymphatic vessels within the epicardium remain to be elucidated.

Recent studies have revealed an underappreciated role of cardiac lymphatic vessels in promoting beneficial effects for different aspects of heart pathology, including cardiac edema, hypertrophic cardiomyopathy and scarring^23,24,33–35^. It remains unclear whether adult heart diseases might have developmental causes in cardiac lymphatic defects and difference in cardiac lymphatic vessels might contribute to different responses after myocardial infarction similarly to collateral coronary arteries^36^. To gain more molecular insight of cardiac lymphatic vessel development in human, we performed single nuclei (sn) Multiomic analyses. The concurrent detection of mRNA and chromatin accessibility helps reveal cellular heterogeneity of LECs that reside in the epicardium. We found VEGFC is highly expressed in arterial endothelial cells, providing a molecular basis of the arterial association of cardiac lymphatic development. Furthermore, we identified a novel population of cardiac lymphatic endothelial cells marked by the PROX1 transcriptional factor, the lymphangiocrine RELN, and enriched in binding motifs of ETV transcription factors. Our data also provide a valuable resource to understand cardiac lymphatic vessel development in human and this might form a basis for future understanding of congenital and adult heart diseases.

## Results

### Lymphatic vessels in human hearts are mainly associated with coronary arteries

The association between the lymphatic vessels and the arteries or veins has been well documented for distinct organs^25^. To identify the potential progenitors of cardiac lymphatic vessels for human heart development, we examined the fetal heart samples donated for research. The two main coronary artery (CA), right CA (RCA) and the circumflex artery (CXA), develop before the 9 post-conception weeks (PCW) although the ACTA2+ vascular smooth muscle cells (VSMC) recruitment is gradual and continues during the fetal period together with the vascular plexus expansion^37^. CAs at this stage have already acquired their arterial identity, as shown by the presence of endothelial cells positive for Notch1 activity and surrounded by the VSMCs (Fig.1A).

**Figure 1.**
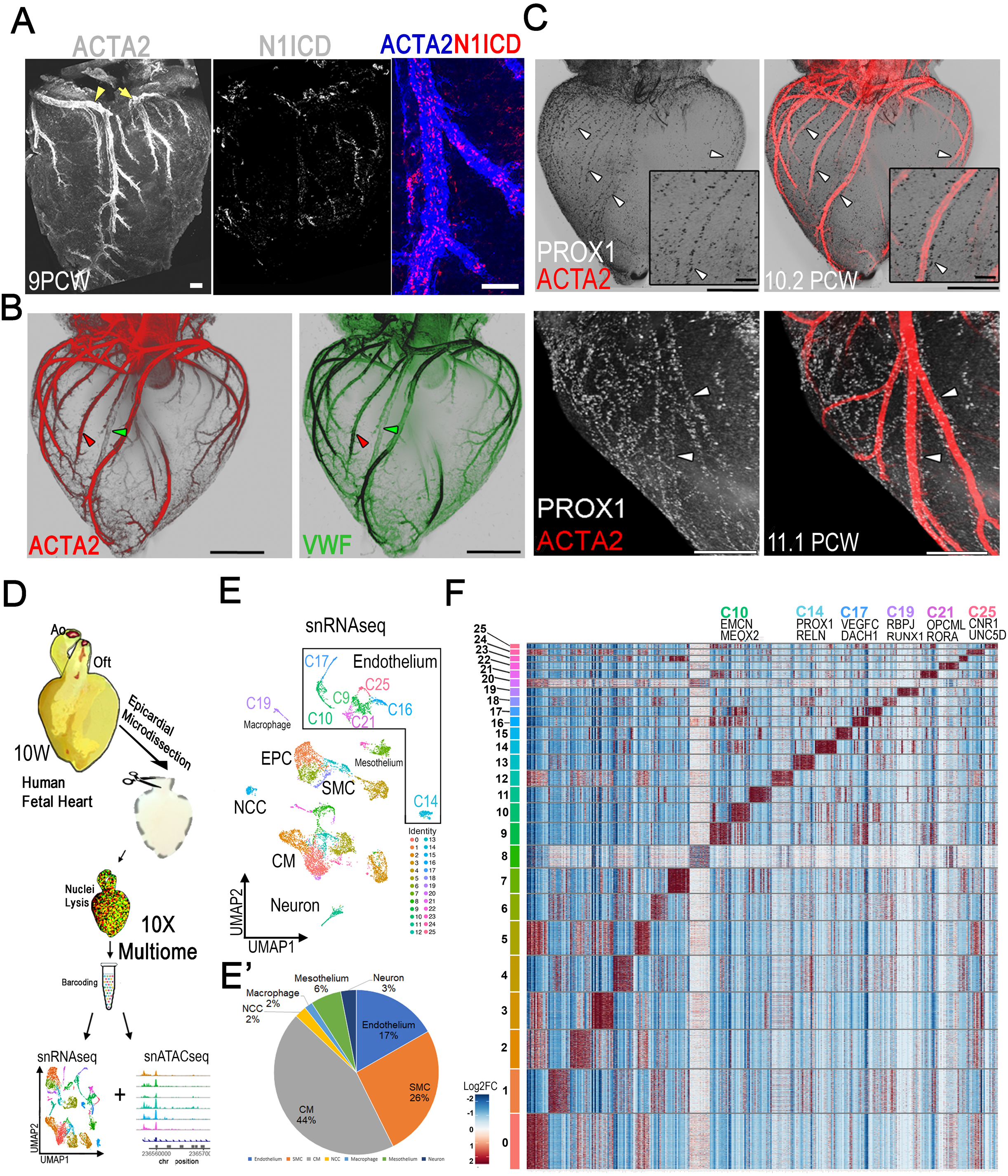
Single nuclei multiomic analysis of human cardiac lymphatic vessels that are mainly associated with coronary arteries. (A) Whole mount dorsal view of 9PCW human fetal hearts immunostained for ACTA2 and N1ICD. The RCA (arrowhead) and the CXA (arrow) are marked (n=3). (B, C) Whole-mount light sheet images of the 10.2PCW and 11PCW (bottom panels) dorsal view of human fetal hearts stained for smooth muscle marker ACTA2 (red) and endothelial cell marker VMF (green) (B) and lymphatic endothelial cell-expressing PROX1 (grey) (C). Magnified views (inset) show details of the RCA. Red arrowheads in (B): CA; Green arrowheads in (B): CV. White arrowheads in (C) indicate the PROX1+ cell nuclei marking LECs associated with the ACTA2+ arteries on the ventricle (n=15). PROX1+ nuclei (lymphatic) primarily align along artery, but also rarely with vein. (D) Schematic of the isolation and enrichment of epicardium before the cell dissociation for the 10X snMultiome profiling. (E) UMAP of snRNA-Seq data for all cells of the 10 PCW heart showing several endothelial clusters (E’) Pie chart showing percentage of all cell clusters from the snMultiome analyses (F) Heatmap of the top 15 differentially expressed genes (DEGs) by Log.fold change for each cluster compared to the rest of the dataset at 10PCW. Scale bars, 200 μm.

We found the arterial identity of the ECs in coronary artery correlates with the onset of blood flow in human heart temporal-spatially (Fig.1A). At 10 PCW the RCA and the CXA are covered by ACTA2+ VSMC cells both in the proximal and in the distal segments while the veins show weak or no ACTA2 staining (Fig.1B). PROX1, a known master homeobox transcriptional factor required for lymphatic differentiation^38,39^ is expressed in human lymphatic endothelial cells (LECs) that follow the course of the main branches of the CA at 10 PCW and acquire a tubular structure (Fig.1C). The outer layer of the ventricle is covered more extensively by these lymphatic vessels associated with coronary arteries by 11 PCW (Fig.1C) although association with some coronary veins are also observed.

### Single Cell Transcriptomic Analysis revealed subpopulations of cardiac endothelial cells

To identify potential endothelial populations that might account for the different associations with arteries or veins, we used a snMultiomic approach to determine the transcriptomics and gene regulatory landscape of human fetal cardiac ECs. This analysis integrates data of snRNA-seq and snATAC-seq collected from the same cells and can enhance resolution of the cell clusters. Cardiac lymphatics were enriched by micro-dissecting the epicardial layer of the heart surface (Fig.1D). After quality control filtering using Cell Ranger (Table S0),10,652 nuclei were processed and 10,610 were plotted for snRNAseq (Fig.1E, Table S1E, S1_2). Data analyses were performed as described in the Online Method, and processed data will be submitted to the gene ontology omnibus (GEO). The uniform manifold approximation and projection (UMAP) for the 10PCW snRNA-seq revealed 26 cell clusters in human fetal hearts (Fig.1E and Table S1e). Clusters related to the endothelium, including all arterial, venous and lymphatic endothelia were identified in clusters 9,10,14,16,17,21,25 (Fig.1F). Heatmap showed snRNAseq gene expression data of the top 15 differentially expressed genes (DEGs) by adjusted p-value <0.05 compared to the rest of the dataset at 10PCW and determined reliable markers for each population (Fig.1F, Table S1F and Source Data 1F).

### Integrative analysis of transcriptome of the human lymphatic clusters

The ‘‘Weighted-Nearest Neighbor’’ (WNN) analysis conducted for the multimodal integration of the snRNAseq and snATACseq enhanced the resolution of all the clusters, on both transcriptional and epigenomic level (Fig.2A). The clusters related to the endothelium were further separated into arterial (cluster 17) (Suppl.Fig.1), venous (cluster 10) (Suppl.Fig.2) and lymphatic endothelia. The expression of the arterial identity marker genes *DLL4, SOX17, NOTCH4* was predominantly restricted to the CA endothelium (Supp.Fig.1). Markers genes such as *NOTCH1, KDR*, and *UNC5B* were expressed in both arterial and lymphatic clusters (Supp.Fig.1) while *NRP2, NR2F2* and *FLRT2* were expressed in both venous and lymphatic markers. These results do not support the presence of potential arterial-lymphatic or venous-lymphatic progenitor populations that might account for the preferential association of cardiac lymphatics with CA.

Joint analyses of the RNA levels by snRNAseq and snMultiome with the WNN integration further improved our ability to define and to resolve the cellular states of the LECs and suggested different gene expression between the LECs (PROX1+/FLT4+) and the macrophage clusters (*MRC1+*/*LYVE1+*). *PROX1* and *FLT4* gene expression was the highest in cluster 14 (Fig.2A,B,C). *PDPN, LYVE1* and *MRC1* showed more distinct expression patterns, suggesting non-specificity of expression at this stage (Fig.2B, C). In contrast to the endothelial specific expression of lymphatic markers in cluster 14 that we identified as LECs (Fig.2B and Table S2d), *PDPN* also showed high expression in mesothelium (cluster 7, 23) while *MRC1* and *LYVE1* were both co-expressed in macrophage cluster 19 (Fig.2B,C). Notably, gene ontology analysis of molecular functions for the LECs revealed VEGF signalling and hyaluronic acid binding activities (Fig.2D and Data S2e). The formation of tubular structure in the epicardium by PROX1 marking LECs is likely a consequence of the activation of the VEGF pathway (Fig.2E). Additionally, we validated the expression of KDR/VEGFR2 in the LECs that formed by elongating along the main coronary arteries in the epicardium (Fig.2F,F’).

Immunostaining of N1ICD, PROX1 and ACTA2 confirmed the preferential association of LECs to CA on the human heart of 11PCW (Fig.2G). VEGFC whole mount immunostaining (Fig.2H,H’) revealed high expression around the CA and validated the expression that we predicted *in-silico* by Feature plot (Fig.2I). Interestingly the locus for VEGFC suggest a specific activation of potential enhancers that are found only in the arterial clusters but not in other endothelial or epicardial cells (Fig.2J). In zebrafish embryos, it is shown that Cxcl12 chemokine signals to Cxcr4 receptors expressed in LECs and guide the arterial association during trunk lymphatic vessel development^40^. However, we only found CXCR4 expression in arterial endothelial cells but not in LECs (Fig. 2I), consistent with the finding in cardiac lymphatic development in zebrafish hearts^23^. These data suggest that the high expression of VEGFC, a known factor involved in lymphangiogenesis found in CA endothelium and in pericytes might account for the association of the nascent cardiac lymphatics to the CA in human fetal hearts.

**Figure 2.**
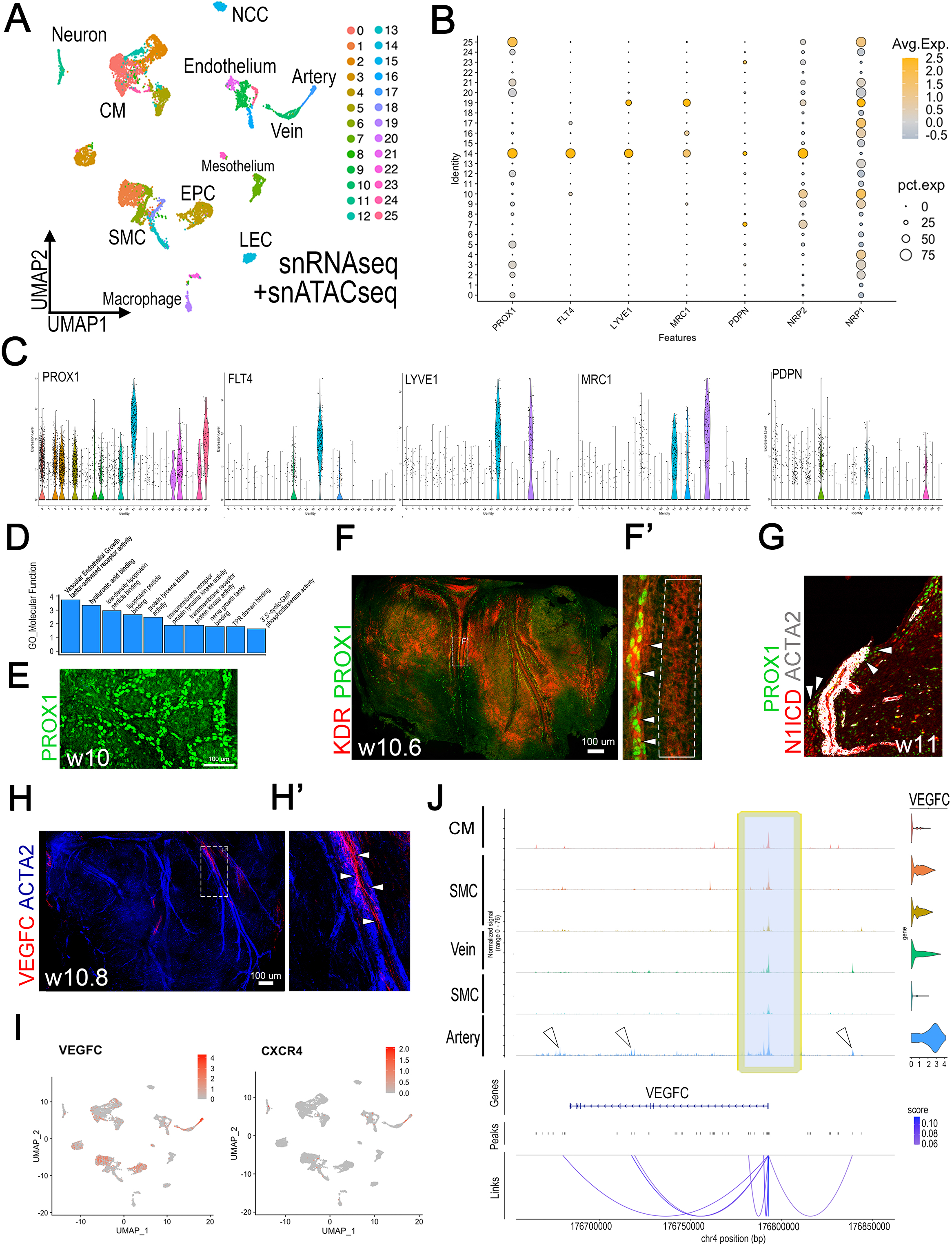
Human cardiac lymphatic endothelial cells are associated with coronary arterial cells that express VEGFC. (A) FeaturePlots of snMultiome (snRNAseq + snATACseq) for the previous lymphatic markers with a higher resolution. The dimension reduction from the WNN analysis shows that cluster 19 *(MRC1/LYVE1)* is close to the LECs of cluster 14 suggesting common open chromatin regions for both populations. (B) Dot plots showing *PROX1,FLT4,LYVE1,MRC1,PDPN,NRP2 and NRP1*. All these markers except for *NRP1*, coexist in Cluster 14 and are expressed in a mixed pattern in the other clusters. (C) Violin plots of lymphatic differentiation markers among different clusters. (D) GO analysis of Molecular Function of the Cluster 14 identify VEGF activity and hyaluronic acid binding as main function. (E) Immunostaining of PROX1 on the dorsal side of human ventricle showing tubular structure formation. (F) Whole mount dorsal view of 10PCW human fetal hearts immunostained for PROX1 (green) and KDR (red) (below). Arrowheads indicate a region where PROX1^+^ cells are enriched in KDR expression (n=3). The dashed box in F shows the zoomed-in area in F’. The dashed box in F’ indicates KDR expression in LECs around CA. (G) Immunohistochemistry for N1ICD (red), PROX1 (green), and ACTA2 (white) on 11PCW human transverse section; the magnified view shows co-staining PROX1+ cells surrounding the CA (white arrowheads) (n=3). (H) Whole mount dorsal view of 10PCW of human fetal hearts immunostained for VEGFC (Red) and ACTA2 (Blue). The magnified view indicates a region where VEGFC is detected (white arrowheads), in close proximity of CA (n=2). (I) FeaturePlots illustrating RNA expression levels from the WNN analysis of VEGFC (left) and CXCR4 (right) are high in arterial endothelium. (J) Pseudobulk of the genome accessibility for the VEGFC locus for selected clusters from the snMultiome and the correlation coefficient between the accessibility of the peak and RNA expression of the gene showing positive correlation between the potential enhancers (white arrowheads) and the promoter of the VEGFC gene in artery. Scale bars, 100 μm.

### Chromatin profiling and genomic organization of the Human Cardiac LEC

Following the arteriovenous endothelial differentiation controlled by NOTCH and NR2F2/COUP-TFII transcription factors, SOX18 starts to activate PROX1. Consequently, the cooperation between NR2F2 and PROX1 trigger the lymphatic differentiation program^41^. To elucidate the mechanisms of LEC heterogeneity, we further examined the chromatin landscape of human fetal hearts. The snATACseq data help resolve the dimensional reduction in the UMAP of the macrophage and the mesothelium populations with the detection of cell-type-specific open chromatin regions (Fig.3A and Table S3A). We used the Multimodal analysis to improve the resolution of all the cell populations before performing the Motif enrichment analysis (Fig.3B). Regions of altered accessibility and gene expression identified ETV2 transcription factors and other ETS variant transcription factor ETS1 and ETS2 (Fig.3C). The ETS family of transcription factors is known to play important roles in embryonic haematopoiesis, vasculogenesis and angiogenesis^42^. Interestingly, Etv2, an endothelial specific and highly evolutionarily conserved ETS transcription factor essential for vascular differentiation also regulates lymphangiogenesis in zebrafish by directly promoting the expression of *flt4*/*vegfr3* within the cardinal vein^43^.

**Figure 3.**
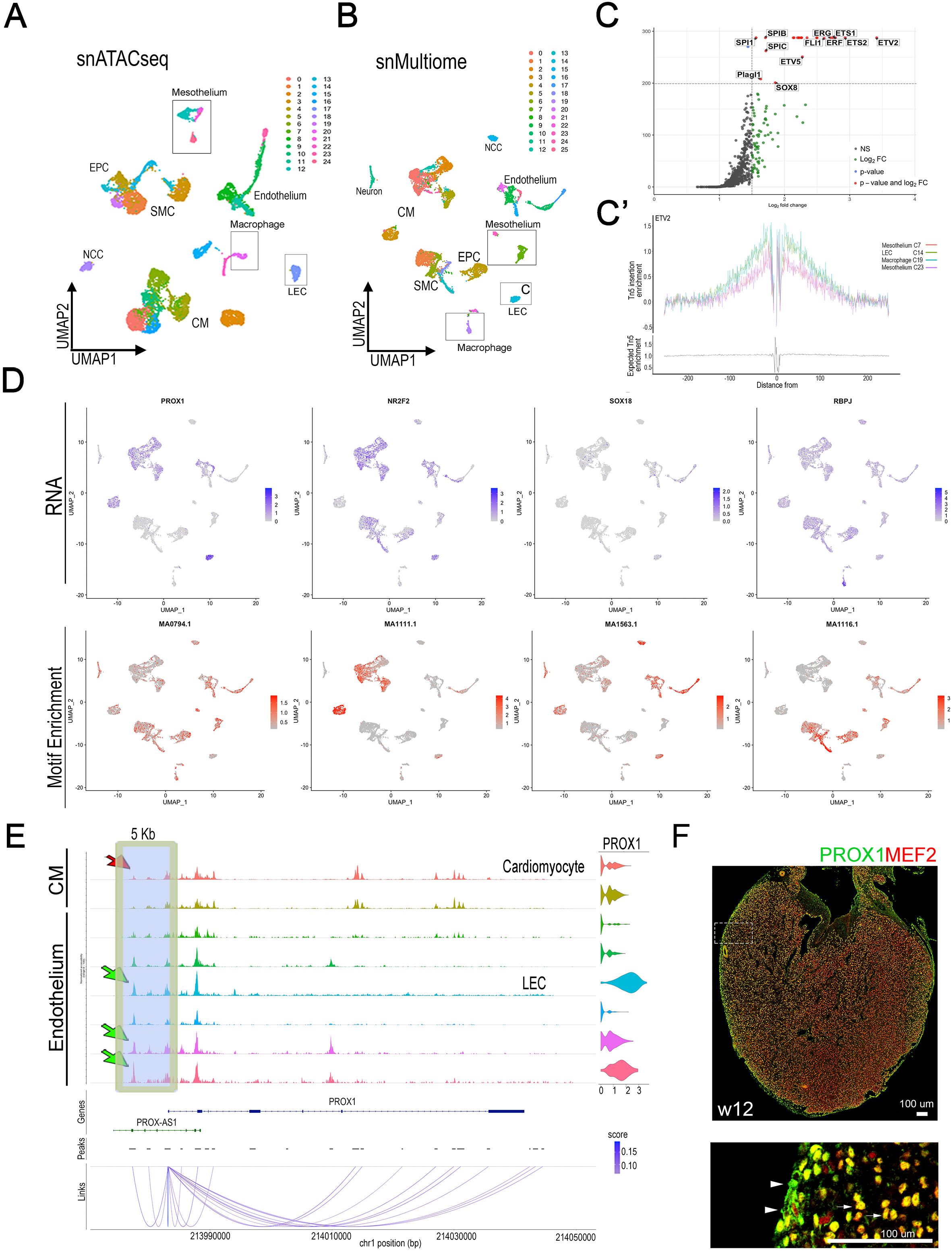
Chromatin profiling of Human Cardiac Lymphatic Endothelial Cells. (A) UMAP of snATACseq and of (B) snMultiome as a result of the WNN integration of snRNAseq and snATACseq datasets. (C) Volcano plot showing positive enrichment of motifs from the LEC cluster 14. (C’) Probability of Tn5 insertion across the genome centered around predicted ETV2 binding motif. The LEC cluster (cluster 14) and in the macrophage cluster (group 19) have a higher enrichment compared to the mesothelial cells (group 7,23) (D) FeaturePlots illustrating RNA expression (upper panels) from the WNN analysis of *PROX1, NR2F2, SOX18* and *RBPJ* and FeaturePlots showing the Motif enrichment for PROX1, NR2F2, SOX18 and RBPJ (lower panels). (E) Pseudobulk of the genome accessibility for the PROX1 locus for selected clusters from the Multiome and the correlation coefficient between the accessibility of the peak and RNA expression of the gene showing positive correlation between the 5kb Enhancer and the Promoter of the PROX1 gene. Red arrow: peak in cardiomyocytes, Green arrow: peak in LECs and other PROX1+ ECs. (F)12PCW fetal heart immunohistochemistry for MEF2 (red) and PROX1 co-staining (green) (n=3); the magnified view shows co-staining of MEF2 and PROX1 in cardiomyocytes (white arrows) and PROX1 staining epicardial LEC (arrowheads). Scale bars, 100 μm.

However, it is still unknown if ETV2 has a direct regulation of PROX1 expression in lymphangioblasts. The presence of motif enrichments for ETS1, ETS2 and ETV2 transcriptional factors in the cluster 14 suggests a potential progenitor nature of this population similar to the venous endothelium. We further examined enrichment of Tn5 insertion integration events nearby the ETV2 binding motif sites by carrying out TF footprinting, delineating a stronger enrichment of integration events flanking the binding motif in the LECs and macrophage groups 200bp upstream and downstream compared to the mesothelium groups (Fig.3C’). The cluster 14 also showed low expression of SOX18 and the NOTCH signaling effector, RBPJ, but medium to high expression of NR2F2 and PROX1, respectively. Interestingly, cluster 14 showed high SOX18 (MA1563.1) but low in PROX1 binding motif (MA0794.1) (Fig.3D). It has been shown that high levels of PROX1 expression direct lymphatic differentiation; and the activity of Prox1 is necessary and sufficient to acquire a lymphatic endothelial cell (LEC) fate by repressing the blood endothelial specification cascade^44^. Therefore, the low enrichment of PROX1 binding motif and high enrichment of ETV2 motif in the lymphatic cluster 14 suggests the levels of ETV2 activation in these cells might prevent the differentiation of lymphatic endothelium, further suggesting a progenitor-like phenotype. These results suggest that transient activation of ETV2 gene expression might be important for the upstream regulation of lymphatic differentiation. Furthermore, this discrepancy between the gene expression and motif enrichment for the transcriptional factors PROX1, NR2F2, SOX18 and RBPJ further suggests a more complex regulation of the lymphatic differentiation (Fig.3D and Suppl.Fig.3D).

Motif footprinting for the PROX1 binding motif in those clusters show high Tn5 insertion 100bp downstream of the motif, suggesting a potential nucleosomal relationship (Suppl.Fig.3E). Likewise, NR2F2, RBPJ and SOX18 motif footprint in the same populations demonstrate positive enrichment of Tn5 insertion genome-wide ±100 bp around their predicted motifs (Suppl.Fig.3E). The top enrichment motifs identified in the LECs revealed a unique chromatin profile (Fig.3C,C’ and Table S3C), different from the macrophages and mesothelium (Extended data Suppl.Fig.4B, 4C), suggesting that they have potentially distinct genomic regulation. Interestingly, comparison of shared motif enrichment between LECs and macrophage clusters but not the mesothelium cluster identified SPIC and SPIB, two ETS transcription factors, respectively involved in lymphoid and leukaemia and ETV6 along with other ETV paralogs also involved in hematopoiesis and maintenance of the developing vascular network (Suppl.Fig.4D,D’ and Extended data of Fig.S4D).

The LEC cluster showed high enrichment for ETS transcription factors binding motif concurrent to low level of the transcripts. We found that the dynamics of transcriptional regulatory mechanisms controlling ETS genes diverge from their RNA expression and therefore their binding activities (Fig.3C, Suppl.Fig.4E). ETV2, ETV5 and ETS2 were present at very low levels or absent in LECs. Only increased levels of ETS1 transcripts (Suppl.Fig.4F) correlated with enriched binding motifs. Considering the complex regulation by transcription factors, we investigated the locus of PROX1 to validate our datasets using the known regulatory elements of the lymphatic genes. We first examined the enhancer region 5kb upstream of the *PROX1* promoter where conserved NR2F2 binding sites were identified^45^.

Consistent with this finding, the pseudo bulk ATACseq analysis of the *PROX1* genomic locus showed an open chromatin state in cluster 14, and in other EC clusters but not in cardiomyocyte clusters (Fig.3E). Co-immunostaining for PROX1 and MEF2 validate the broad expression of PROX1 in cardiomyocyte and suggests that only higher levels of PROX1 (peak in its enhancers found in LECs) are able to induce the specific lymphatic differentiation cascade (Fig.3E,F).

### PROX1, RELN, and ETV binding motifs define the human cardiac LECs

To further examine EC heterogeneity and compare LECs to other ECs, we generated a UMAP organized in 7 clusters (Fig.4A, B). The integrative analysis conducted on all endothelium by hierarchical clustering of joint snRNAs and snATACseq helped identify a unique cluster among all others that showed highly similar gene expression and gene activity for, *PROX1, NRP2, RELN, STAB2, STON2, VAV3, CD36* thus defining a *bona fide* cardiac LEC population with a very confined expression of ETV2 (Fig.4C). Furthermore, we validated RELN expression by immunostaining using an antibody against the secreted form of RELN. We found RELN expression nearby all the PROX1+ LECs suggesting a potentially evolutionarily conserved role of this lymphangiocrine (Fig.4D). Spearman correlations analyses revealed significant correlations between RNA levels of PROX1 and ETV2 and between the binding motifs of NR2F2 and ETV2 (Fig.4E).

**Figure 4.**
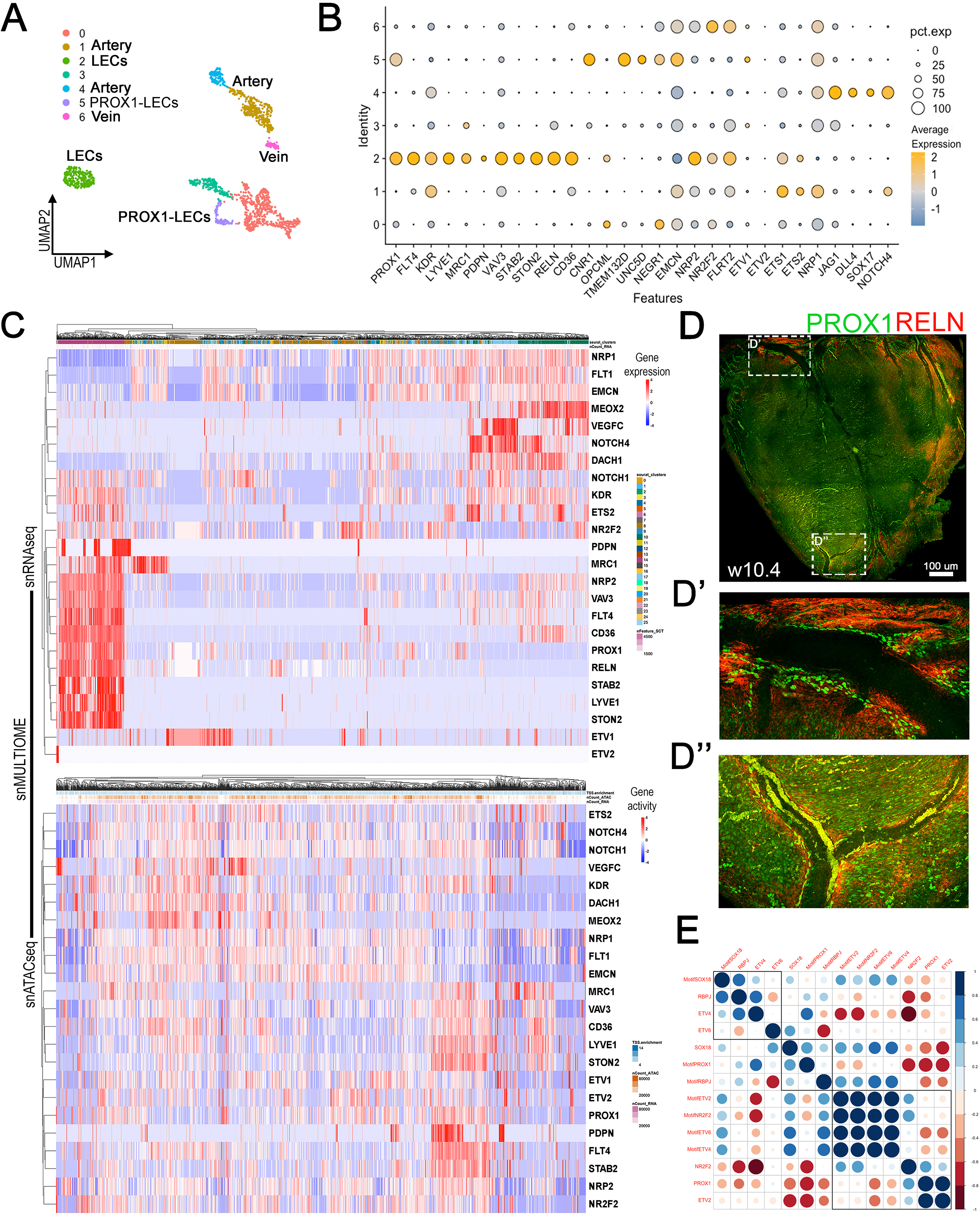
A *bona fide* population of cardiac lymphatic endothelial cells identified in human fetal heart. (A) UMAP generated by EC subclusters of the cell dataset. (B) Dotplot depicting both the expression level and the percentage of cells expressing selected genes from the EC clusters and lymphatic markers. (C) Heatmap generated from the EC clusters and depicting gene expression from snRNAseq and of the snATACseq showing genes involved in lymphatic cluster. (D) Whole mount dorsal view z-projection of 10PCW human fetal hearts immunostained for PROX1(green) and secreted form of RELN (red) (n=2). Insets (D’ and D’’) show regions of the ventricle, including the apex where PROX1+ cells are secreting RELN. (E) Spearman correlation plot for selected TFs average RNA expression and Binding Motifs presence in open chromatin of endothelial and lymphatic cells. Colour intensity of the circle indicates the strength of the correlation according to the scale provided at the side of the graph. Scale bars, 100μm.

### ETS-transcription factors control the LEC heterogeneity

Sub-clustering analysis of the LECs further facilitated the detection of different levels of expression of PROX1 and FLT4/VEGFR3. (Fig.5A,B). This result suggests that LECs are heterogeneous and 4 sub-clusters were identified. We hypothesized that the heterogeneity of the LECs might be depending on other transcriptional factors, acting upstream of the canonical LEC differentiation cascade. To further characterize the heterogeneity among the LECs, we performed motif enrichment analysis of the LEC subclusters 0-3 and we identified ETV1 and ETV4 as the top two motifs (Fig.5C) and allocate the metallopeptidase/ metalloendopeptidase activity as the main GO molecular functions (Fig.5D). Opposite trends in expression levels between ETS Variant Transcription factors (ETV1, ETV2, ETV4, ETV5) or other ETS transcription factors and PROX1, NR2F2, RBPJ were found in the analysis of the LEC subclusters suggesting that there might be an antagonistic relationship between these factors (Fig.5E). Unlike the other ETV factors and similar to PROX1, ETV6 showed very high expression in all the LEC subclusters (Fig.5E,F). The binding activities of these factors in region of open chromatin suggests that they might assist the plasticity of the LECs (Fig. 5G).

**Figure 5.**
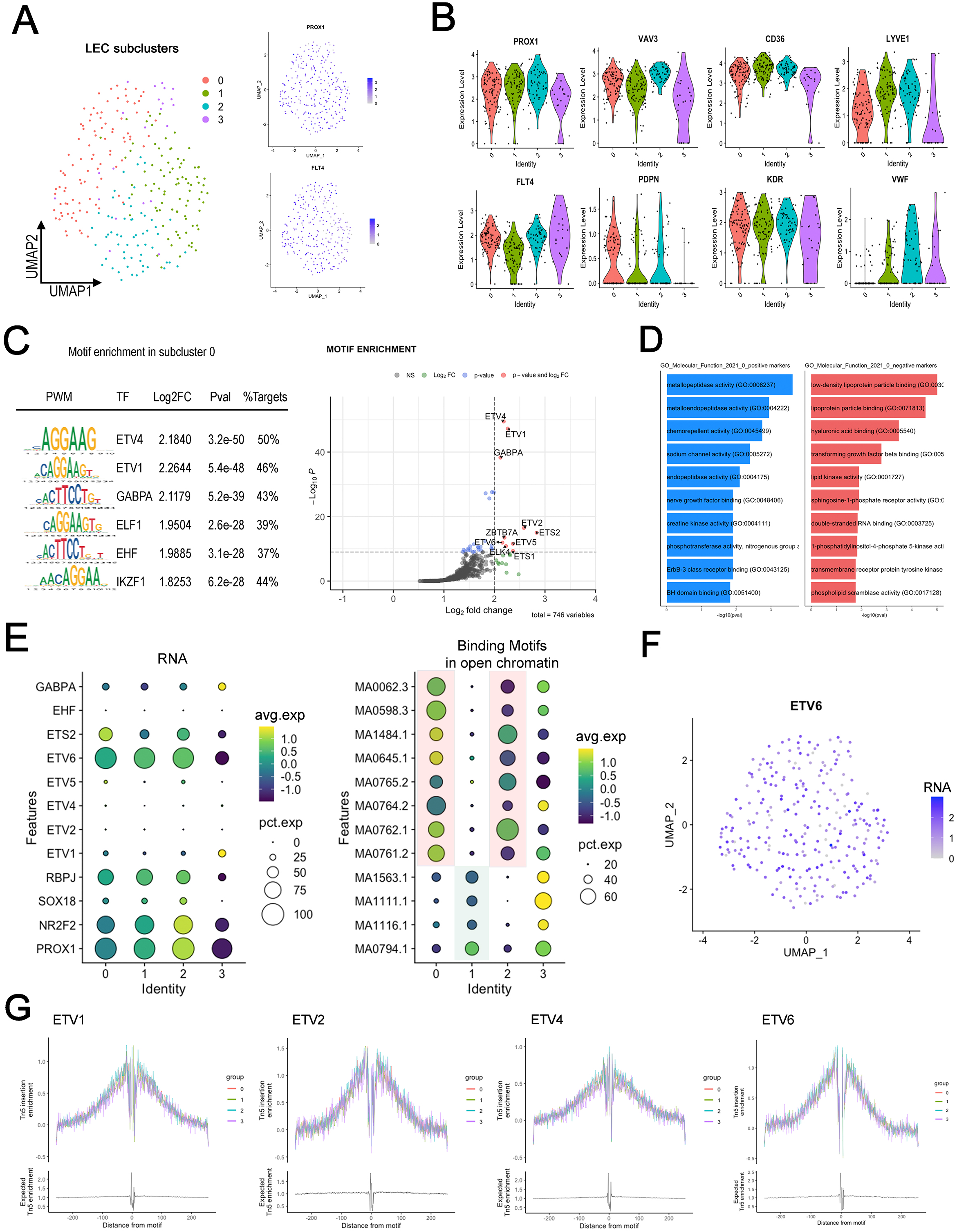
Subcluster analysis of LEC heterogeneity. (A) UMAP generated by subclustering the LECs from cluster 14 (left). FeaturePlots of snRNAseq + snATACseq from the subclusters of the LECs showing different levels of expression of PROX1 and FLT4 RNA (right). (B) Violin plot depicting the expression levels of lymphatic markers and endothelial markers among the four subclusters. (C)Top 6 enrichment motifs for the subcluster 0 compared to the rest of the LECs (left), and Volcano plot showing positive enrichment in the subcluster 0 of all the ETS transcriptional factors (right). (D) GO term analysis of the molecular function predict positive enrichment of metallopeptidase and metalloendopeptidase activity. (E) Dot plots illustrating RNA (left) and binding motifs (right). High level of expression for PROX1, NR2F2 and RBPJ and ETV6 and ETS2 only. Binding motifs for all the ETV family members show trends opposite to PROX1, NR2F2, SOX18 and RBPJ in the LEC subclusters. (F) FeaturePlots of the subclusters in the LECs showing high levels of ETV6 expression from all the cells. (G) Normalized probability of Tn5 insertion across the genome centered around predicted ETV1, ETV2, ETV4 and ETV6 binding motifs comparing the 4 subclusters.

We further performed Differentially Accessible Regions (DARs) analysis on the entire datasets to identify novel predicted enhancers in the genes involved in lymphatic differentiation (Fig.6A,B and Table S6A). First intron after the first exon of *FLT4* gene and the 5’ intronic region upstream of the *STAB2* promoter were enlisted as the top DE peaks (Fig.6B,C,C’,D,E’and Table S6B). As shown originally by the SHARE-seq, simultaneous high-throughput ATAC and RNA expression can be used to correlate chromatin-gene expression^46^.

**Figure 6.**
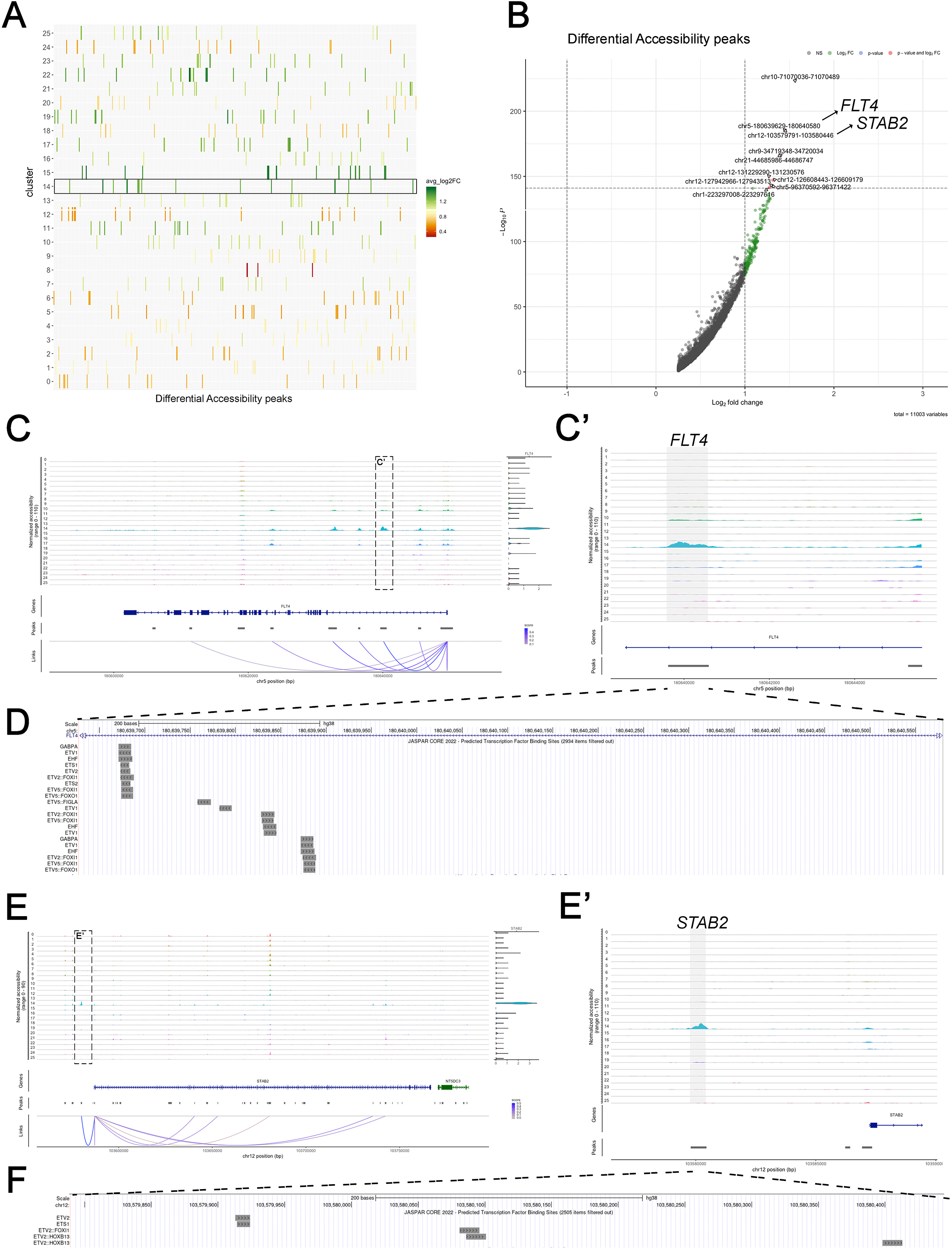
Predicted enhancers from differentially accessible chromatin peaks. (A) Plot showing avgLog2FC of the top 15 differentially accessible (DA) peaks from all the clusters compared to the entire dataset (B) Volcano plot showing positive enrichments of DA peaks position from the LEC cluster (C) Pseudobulk of the genome accessibility for the *FLT4* locus from the snMultiome showing positive correlation between the accessibility of the peak and RNA expression of the gene (Link-Peaks to Gene). (C’) Magnified view of boxed area of the Pseudobulk showing the first intron from *FLT4* locus. (D) UCSC Genome Browser view of the JASPAR2020 predicts a high number of ETS transcriptional factors binding motifs. (E) Pseudobulk of the genome accessibility for the *STAB2* locus from the snMultiome showing positive correlation between the accessibility of the peak and RNA expression of the gene (Link-Peaks to Gene). (E’) Magnified view of boxed area of the Pseudobulk showing the intronic region upstream of*STAB2* promoter. (F) UCSC Genome Browser view of the JASPAR2020 predicts binding site for the ETS transcription factors.

We applied the link peaks to gene function of Signac to compute the Pearson correlation for the accessibility of the STAB2 and FLT4 peaks to their gene TSS, corrected by GC content, overall accessibility, and peak size. The top 0.5 scores link these two noncoding DNA elements to the expression of their nearby genes (Fig.6C,E). JASPAR2020 predicted binding motifs from these two peaks identified several binding motifs for the ETS transcriptional factors, especially in the FLT4 locus (Fig.6D,F). None of the PROX1/NR2F2/SOX18/RBPJ motifs were found in these chromatin regions. Altogether, this data confirms the validity of the datasets and identifies two novel cis-acting DNA elements large ∼1kb involved in the regulation of the LEC differentiation. These two enhancers are regulated by the ETS transcriptional factors and are independent of the canonical LEC differentiation cascade operated by PROX1. More studies are required to test if these genomic regulatory elements are sufficient to induce the lymphatic differentiation cascade and how deep its conservation is across animal kingdom.

## Discussion

Here we report the first snMultiomic analysis of the development of the cardiac lymphatic plexus of human fetal hearts. The snMultiomic analyses allowed to simultaneously define the gene expression profile and open chromatin landscape from these lymphatic cells using the same nuclei. This reduces potential variability caused by performing chromatin and gene expression analyses in separate assays. We have successfully created a cellular and molecular atlas of human fetal cardiac lymphatic vessels that develop along coronary arteries, providing transcriptomic and epigenomic map of the early steps of lymphangiogenesis. Our integrative analysis showed that the subepicardial plexus consists of multiple heterogeneous populations of ECs and LECs. We found that coronary arterial endothelial cells express high levels of VEGFC, which might account for the arterial association of cardiac lymphatics. These results suggest that the preferential association with CAs might not be due to the physiological differences between arterial and venous functions; instead, it depends on which tissues express VEGFC. This might help explain the different associations observed in zebrafish, mice, and human fetal heart and different organs.

Our results also support prior research showing the LEC fate is directed by PROX1 as the key molecule marker for the majority of cardiac LECs and regulated by the equilibrium among the cell fate regulators NOTCH1^47^ and NR2F2^41^, and that lymphatic plexus formation follows the VEGFC/VEGFR3 cascade. Our data indicate that the promoter of PROX1 has several degrees of activation that are tissue specific. The high transcriptional level of PROX1 is a necessary step to drive LEC fate specification and relies, conjointly on the regulation of factors bound to the well-known element found 5kb upstream of the TSS, also on other enhancers that need to be identified. The regulatory region upstream of PROX1 shows activation in several endothelial clusters, each of them showing different levels of gene transcription (Fig.4A), suggesting that the activation of this enhancer might be part of a larger genomic regulation. This complex chromatin organization in LEC suggests the differentiation is a very plastic process that may involve more epigenetic marks to sustain intermediate steps.

We validated the presence of a recent identified lymphoangiocrine molecule known as RELN^48^ in the human cardiac LECs. Our data help identify a novel population of LECs in the epicardium. However, further studies are required to fully characterize the origin and the differentiation potential of the PROX1/NRP2/RELN LEC cluster (Table S2d). The expression of VAV3 and CD36 in this population supports the conclusion that these LECs might be from mesenchymal/hemogenic precursors rather than derived from a venous endothelial expansion. Future investigations might need to address the specific location of these cells and to evaluate their potential presence in adult cardiac tissue. Moreover, we have characterized a “macrophage-like” cell cluster that shares a similar genetic differentiation program to LECs and a mesothelial cell population with a distinctive molecular feature. Additional studies are required to test the specific role of these cells in regulating homeostasis of and/or interactions with the cardiac lymphatic vessels and to define the specific roles during cardiac development and disease as they might have a potential compensatory effect in the absence of one or more sources during the expansion of the lymphatic plexus.

The snMultiome data of the human fetal hearts reveal the plasticity and the heterogeneity of the cardiac lymphatic endothelium. The molecular mechanisms that regulate lymphangiogenesis in mammals are conserved in zebrafish where the posterior cardinal vein (PCV) is the origin of the LEC^49^. In mouse embryos, cardiac lymphatics are mainly associated with coronary veins; and the origin of cardiac lymphatics has been related to extra-cardiac venous endothelium with a smaller contribution of a 20% from hemogenic Vav1^+^ non-endothelial progenitors^50^. Furthermore, recent studies in zebrafish has suggested that the outflow tract (OFT) lymphatics are derived from facial lymphatics with addition of venous and non-venous progenitors^24,51^, suggesting that the main contribution might be from specialized angioblasts^52^. Moreover, isolated lymphatic clusters have been identified in zebrafish and mouse hearts^24^. These results suggest that the progenitor sources in human fetal hearts require further investigations.

Our results propose a novel role in the regulation of cardiac LECs by different thresholds of ETV2, as recently reported for endothelial and erythropoietic development in mammals^53^. The potentially transient interaction between ETV2-PROX1 suggests the existence of newly specialized angioblasts and reveals the presence of a novel population during human cardiac lymphatic development similar to the Etv2+Prox1+ lymphangioblasts discovered recently from the paraxial mesoderm of mouse embryos^54^. Complementary to the role of ETV2, other ETS transcription factors, such as ETV1 or ETV4, antagonize the lymphatic differentiation signalling cascade directed by PROX1, while TEL /ETV6, an evolutionarily conserved ETS repressor, and a known master regulator of hematopoiesis in the bone marrow^55,56^ with essential role in the regulation of endothelial sprouting^57^, cooperate to uphold the progenitor state. The enrichment of ETS transcription factors binding motifs in these cells provides new evidence for an upstream regulation of PROX1 in guiding lymphatic identity. This demonstrates that the main control of the LEC fate acquisition might be at the genomic level. Moreover, it also suggests that the reprogramming from BEC to LEC regulated by PROX1 might require the fine-tuning by other ETS transcription factors.

A recent study identified distinct EC populations isolating CD31+ cells from scRNAseq of the 13/14 PCW human fetal heart^58^. Our approach complements this study by dissecting out the epicardium, successfully enriching high number of LECs. Taken together our data demonstrate that the human cardiac lymphatic development follows the main coronary arteries, consistent with observations in other large mammals and zebrafish and has a unique organ-specific molecular regulation. Our results might lead to future identifications of potential therapeutic biomarkers for the treatment of lymphatic bed abnormalities or cardiac repair. The simultaneous characterization of a *de novo* cardiac LECs provided in this work by profiling mRNA and chromatin accessibility might further shed light on the complex regulation of the human cardiac lymphatics and provide a better understanding of their early steps of commitment.

## Abbreviations and Acronyms

PCW: post-conceptional week
CA: Coronary arteries
CV: Coronary veins
EC: endothelial cells
LEC: lymphatic endothelial cells
UMAP: Uniform manifold approximation and projection
DEG: differentially expressed genes
WNN: Weighted-Nearest Neighbor
TSS: Transcriptional Start Site
PCV: Posterior Cardinal Vein

## Methods

### Human fetal heart

De-identified human fetal tissue was obtained from elective terminations with informed consent with the approval reviewed by CHLA/USC IRB. Gestational age (PCW) was determined using guidelines from the American College of Obstetricians and Gynecologists using a combination of ultrasound and last menstrual measurements. Tissue was not collected in cases where termination of pregnancy was conducted due to an identified foetal or pregnancy abnormality.

### Human samples microdissection

Samples were dissected under a Leica stereoscope in a ice-cold DMEM media, sterile and RNAse free. The hearts were placed in petri dishes and the forceps were used to delicately dissect out the epicardium of the fetal hearts. The forceps were then used to maintain the human fetal tissues in place while carefully peeling the epicardial/subepicardial layers from the ventricle and from the peritruncal region of the Aorta and of the Outflow tract (PA).

### Light sheet Immunofluorescence

Whole-mount immunolabeling was carried out on human fetal hearts following fixation as the described below. Hearts were bleached in Dent’s bleach (DMSO:H2O2:methanol, 1:2.5:40) overnight, rinsed in methanol and then fixed overnight in Dent’s fixative (DMSO:Methanol 1:4) at 4 □C and washed in PBS + 0.1%Tween (PBST) for 3 to 8 hr the next day. Hearts were then incubated primary antibodies (PROX1, ACTA2) in blocking solution (HIHS:DMSO:PBS, 1:4:15) for 3–5 days. After this step, they were washed with PBST for 6 hr. Hearts were then incubated secondary antibodies in blocking solution overnight, then washed in PBST for 5 hr.

### Human Heart Clearing

Human Heart Clearing Immunolabeled human tissue was cleared using an adapted iDISCO method^59^ after mounting in 2% Agarose (in TBE). Briefly, hearts were dehydrated through a methanol series for 6 hours, then in dichloromethane:methanol (2:1) overnight. Hearts were then transferred to 100% dichloromethane (Sigma 270997-12×100ML) for 3 hours and then dibenzyl ether (DBE, Sigma 108014-1KG) overnight. Labeled tissues were stored in DBE for up to 1 week during imaging. Cleared human fetal samples were imaged with a bidirectional triple light sheet microscope (UltraMicroscope II, LaVision BioTec), and acquired images were compiled in Vision4D (Arivis).

### Immunofluorescence on sections

Paraffin sections of human fetal hearts (7 μm) were permeabilized in PBS 1X + 0.3%-0.5% TritonX100. Antigen retrieval was performed in Citrate buffer pH6 for max ∼10 min and peroxidase quenching on was done for 1hr in 0.3% of H2O2 in Methanol. The sections were incubated overnight with primary antibodies at 4 °C, followed by 1 hr incubation with a fluorescent-dye–conjugated secondary antibody. N1ICD or PROX1 were immunostained using tyramide signal amplification (TSA).

### Whole-mount immunofluorescence

Human fetal heart were fixed for O/N in 4% paraformaldehyde (PFA) in PBS at 4°C. The following day 2 washes of PBS were made. Dissected hearts were prepare for the flat-mount as previously described^60^. Antigen retrieval was performed in Citrate buffer pH6 for max ∼15 min, adjusting the time for specifying antibodies. Peroxidase quenching on dissected hearts were done for 1hr in 0.3% of H2O2 in Methanol. The tissue was then permeabilized for 2 hr with 0.5% Triton X-100/PBS and subsequently blocked for 1day in Histoblock solution (5% goat serum, 3% BSA, 0.3% Tween-20 in PBS). After several washes in PBS-T (PBS containing 0.1% Tween-20), hearts were mounted in VECTASHIELD Antifade Mounting Medium. Whole-mount hearts were imaged with LEICA STELLARIS confocal microscope. Z-stacks were captured every 5 μm. Hearts were incubated with primary N1ICD-antibody at 1:500 for 2 days at 4°C and allowed to settle at RT for 1 hr. This was followed by washing 5 hr in PBS/0.4% TritonX100 at RT. Hearts were then incubated in presence of anti-rabbit HRP (1:500), DAPI (1:1500) and anti-VE cadherin (1:500) overnight at 4°C. After resting for 1 hr at RT, hearts were washed in PBT, and 30 min in presence TSA (tyramides) diluted 1:200 in PBS. The main coronary trees were obtained by Z-projection.

Antibody list for Light sheet: Sigma Monoclonal Mouse anti-ACTA2 -Cy3 1:400, C6198-0.2ML; (1:400); Dako Monoclonal Mouse anti-vWF Clone F8/86, MO616 (1:200); EMDMillipore Polyclonal Rabbit anti-PROX1 antibody, AB5475 (1:300); Antibody list for whole mount/on sections IF: Cell Signaling Technology Monoclonal Rabbit of Cleaved Notch1 (Val1744) Antibody cat.4147 D3B8 (1:150); R&D Goat Polyclonal Anti-Human Prox1 Antibody Cat. AF2727 (1:200) ; Agilent(Dako) Monoclonal Mouse Anti-Human Smooth Muscle Actin, Clone 1A4, M0851 (1:200); EMDMillipore Mouse Monoclonal Anti-Reelin Antibody, a.a. 164-189 secreted reelin, cat. MAB5366 (1:200); Biolegend Mouse IgG1, Monoclonal anti-human CD309 (VEGFR2) Antibody, cat. 359903 (1:200); Novus Biologicals Rabbit Polyclonal anti-VEGFC antibody, cat. NB110-61022 (1:200).

### 10X Multiome Nuclei isolation

Samples were dissected and resuspended in cold DMEM + 0.02% BSA + 10% FBS. Epicardial preparation was first resuspended in WB (Wash Buffer) in freshly prepare, ice-cold lysis buffer (LB). WB: 10 mM Tris pH 7.4, 10 mM NaCl, 3 mM MgCl2. LB: WB + 0.1 % Tween-20 + 0.1% NP-40, 0.01% digitonin + 2 U/mL RNase inhibitor (Roche). Epicardial preparations (2 biological replicates at 10PWC) were directly lysed and high-purity nuclei from live tissue extracted without prior pre-processing for cell dissociation (scRNA) to avoid stress resulting in change in RNA transcripts. Nuclei were incubated for 5 minutes on ice in 100 uL LB. LB was diluted by adding 900 uL of WB, then samples were centrifuged at 500 x g for 5 minutes at 0 C. Nuclei were checked for complete lysis, counted and then resuspended in 1X Nuclei Dilution Buffer (10X), as suggested by the guideline for optimal performance, to a concentration of 3,080-7,700 nuclei/ul in order to target 10000 nuclei Recovery. Samples were after loaded onto the Chromium 10X controller. Libraries were constructed using the 10X Genomics ATAC and RNA Multiome Kit (10x Genomics Chromium Single Cell Multiome ATAC + Gene Expression) immediately after the resuspension from the CHLA SC2 core.

### Library strategy: Multiome (ATAC and RNA)

Raw sequencing data were converted to fastq format cellranger-arc-2.0.0. QC Metrics are provided.

### Data analyses

Multiome analysis was performed as previously described^61^.

### QC and cell filtering

We filtered the analysis based on the nucleosome signal score and TSS enrichment score for each cell. Cells with a nuclear TSS enrichment score >1, a nucleosome signal score <2, between 5,000 and 70,000 total ATAC-seq counts (based on the 10x Cellranger ATAC-seq count matrix) and between 1,000 and 25,000 total RNA counts were selected.

### Gene expression data pre-processing and cell annotation

Gene expression normalization of the UMI count data was performed using SCTransform and PCA obtained from the SCTransform Pearson residual matrix using the RunPCA function in Seurat. We found the 20 nearest neighbors for each cell using the FindNeighbors function, with dims = 1:50 to use the first 50 principal components and annotated cell types in our scATACseq (EpicardialMultiome) dataset by label transfer from our reference dataset (scRNAseq).

### DNA accessibility data processing

ATAC-seq peaks in the Epicardial dataset were identified using MACS2. Peak calling algorithm was run on the fragment file and was performed for each cell type using the CallPeaks function in Signac. Dimension reduction was performed on the DNA accessibility assay dataset using LSI and UMAP as described above. The function FindNeighbors with dimensions = 2:40 and reduction = ‘lsi’ was followed by FindClusters with algorithm = 3.

### Multimodal label transfer

scATAC-seq and scRNA-seq datasets were analyzed separately and considered as independent datasets before the WNN transferring to evaluate the accuracy of the Multiome multimodal transfer procedure. We identified anchor cells between the scATAC-seq and scRNA-seq datasets using canonical correlation analysis, with the function FindTransferAnchors in Seurat with the parameters, reduction = ‘pca’. Cell type labels were transferred from the scRNA-seq to scATAC-seq dataset using the TransferData function, with weight.reduction = query[[‘lsi’]] and dims = 2:30 to weight anchors based on nearest-neighbor distances in the LSI space. To evaluate the accuracy of multimodal label transfer, we counted the number of cells obtaining the correct predicted cell type label.

### Heatmap

dittoHeatmap (dittoSeq Package) function was used for the generation of hierarchical heatmap. Normalized counts in the case of the snRNAseq or gene activity counts in the case of the snATACseq were used to make the Heatmap. The gene activity assay from the scATAC-seq dataset was done by counting fragments overlapping the gene body and a 2-kb upstream region for each gene in each cell, running the GeneActivity function of Signac. The gene activity counts were then log-normalized for the DNA accessibility assay using the NormalizeData function in Seurat.

### Correlation Matrix

The corrplot R package was used to create spearman correlation matrix that automatically reordered and correlate the average expression data and the average motif enrichments from all the ECs clusters.

## Statistical analysis

The data presented are means ± standard deviations. The Mann-Whitney rank-sum test was used for assessing statistical differences between 2 groups,unless otherwise stated as for the Chromatin accessibility and Motif analysis where we used the “LR” test. P values <.05 were considered statistically significant.

## Data Availability

Raw datasets, images and data analyses will be made available upon requests to the corresponding authors.

## Acknowledgements

We thank the Saban SC2 core for the 10X single nuclei Multiome analysis and Cellular Imaging core. We would also like to thank Melissa L. Wilson (Department of Preventive Medicine, University of Southern California) and Family Planning Associates for coordinating fetal tissue collection.

Source of Funding. This study is supported by Saban Core Pilot Award (to ST and CL) and Team Science Research Award (CL).

## Author Contributions

Contribution: ST and MH performed experiments and analyzed data; ST and CL designed research, analyzed data, and wrote the manuscript. MT and BG help provide human samples. MT, BG, TQ, ST, MH and CL edited the manuscript.

