## Supplementary material for "Single nuclei multiomic analyses identify human cardiac lymphatic endothelial cells associated with coronary arteries in the epicardium": Suppl.Fig.

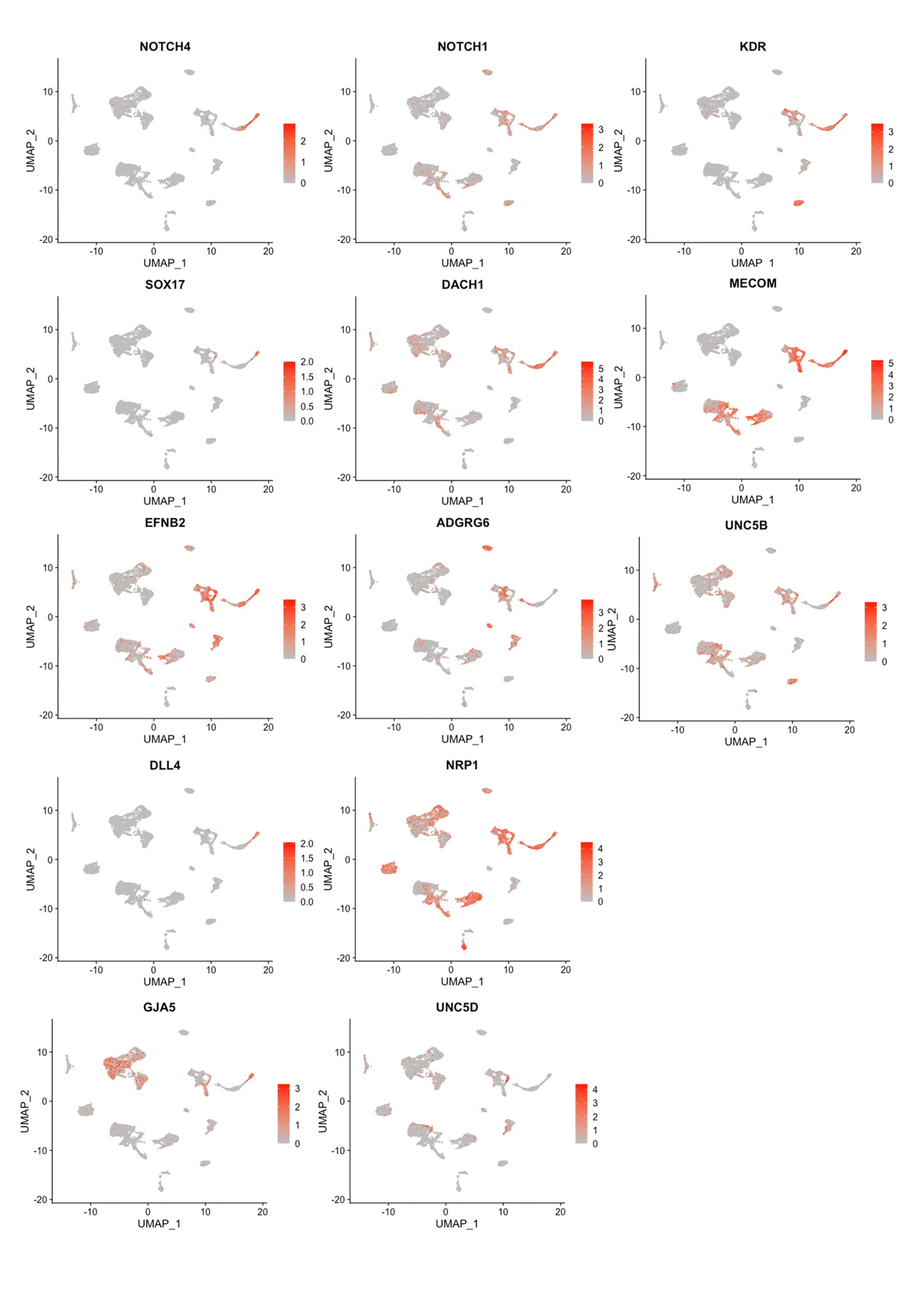
**Suppl. Fig.1. Examples of arterial markers.** UMAP of single-Nuclei multiome data for all cells of the 10 PCW. FeaturePlots depicting high expression for the arterial markers *NOTCH1,NOTCH4,* *KDR1 DACH1 and NRP1* among others*.* *KDR* was highly expressed in the LECs and arterial endothelium. *NRP1* was showing high expression in macrophage cluster and in all the endothelial but not in the LECs.

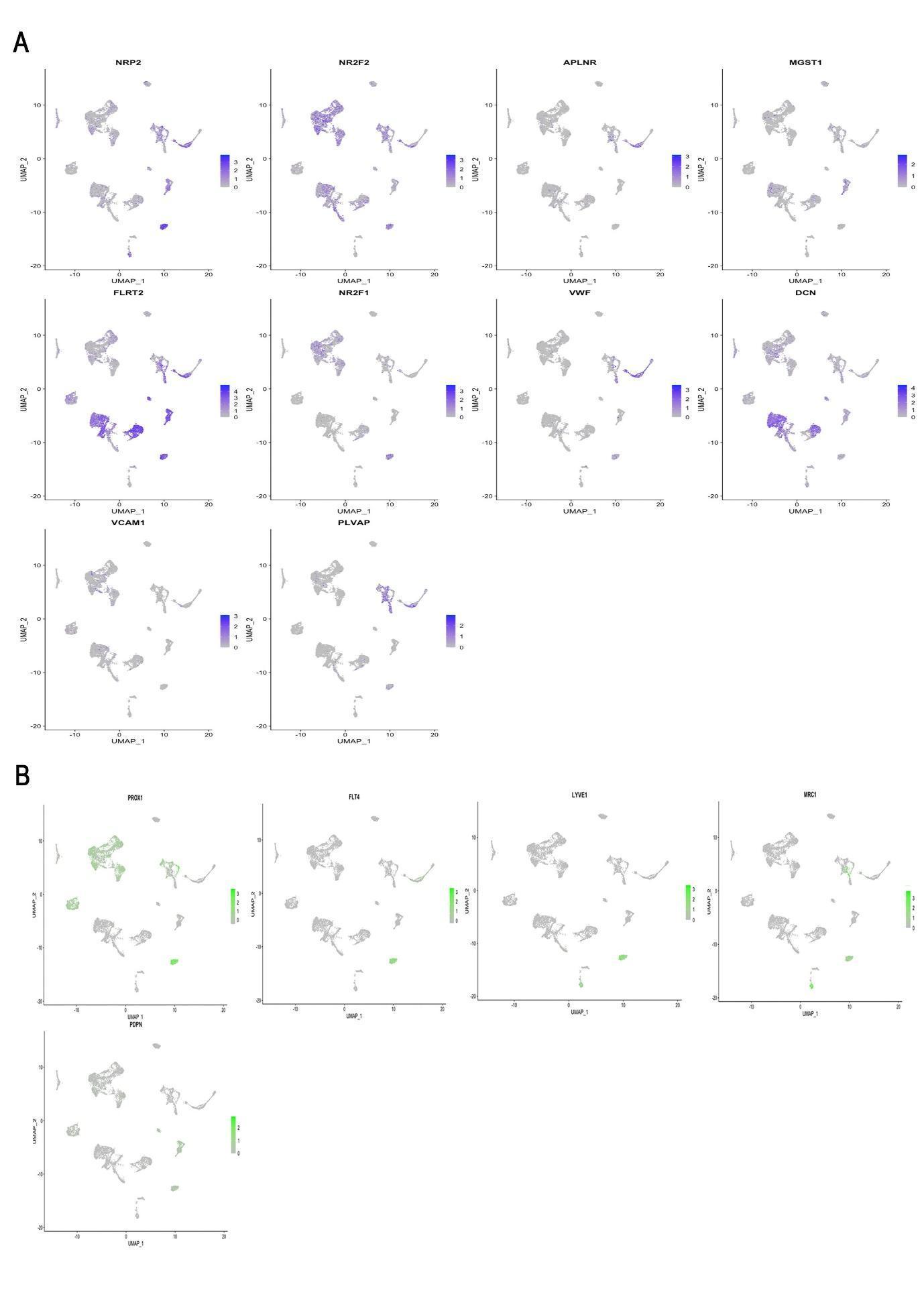
**Suppl. Fig.2. Examples of venous markers** (A) FeaturePlots depicting high endothelial expression for the venous markers *NRP2*, *NR2F2 and APLNR*. *NRP2* shows a significantly high expression in the LEC clusters, and a low expression in the other lymphatic sources whereas *NR2F2* shows a mild RNA expression in several clusters suggesting a prominent role as a transcriptional factor.(B) FeaturePlots depicting high endothelial expression for the lymphatic markers *PROX1*, *FLT4, LYVE1,MRC1* and *PDPN*.

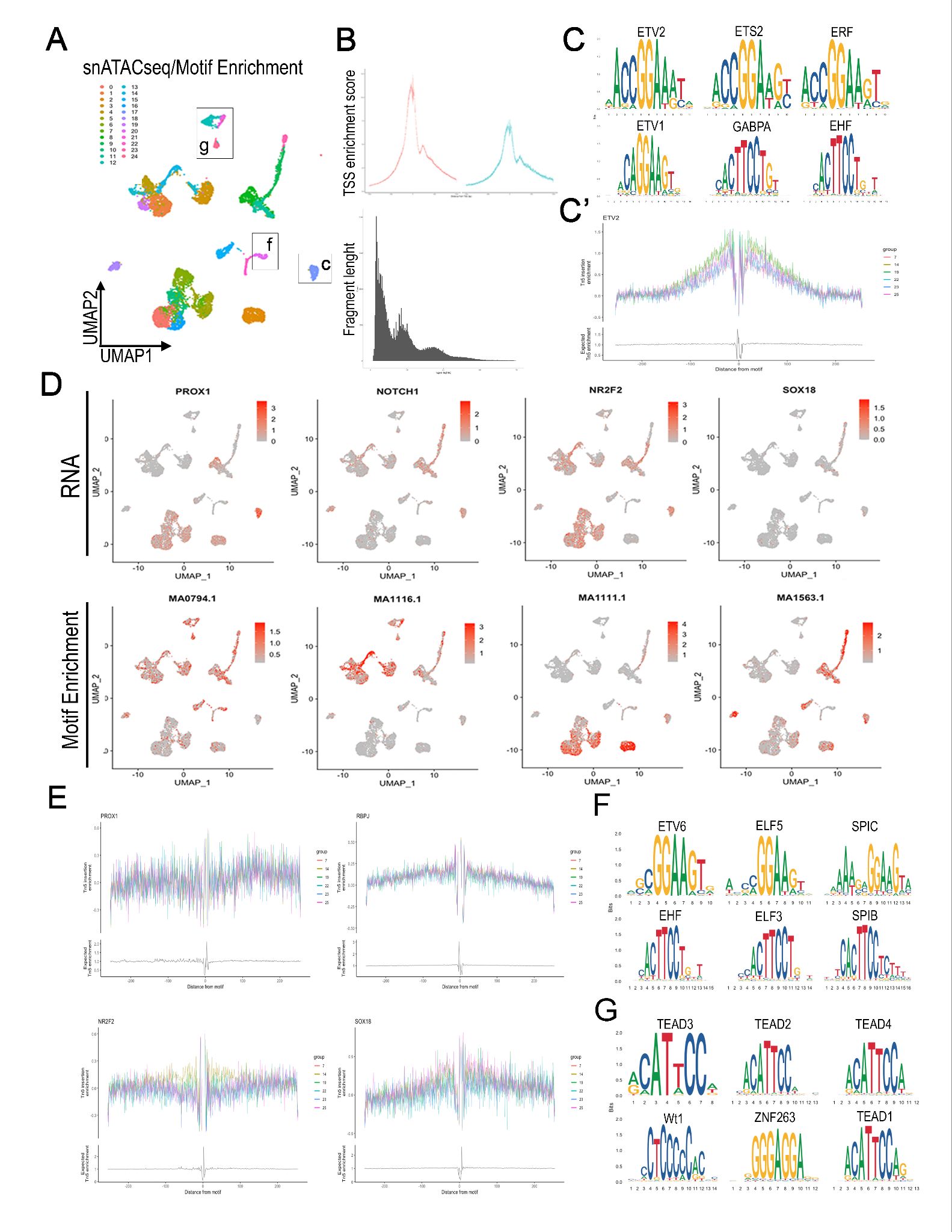
**Suppl. Fig.3 Single-nuclei ATAC analysis of accessible chromatin in LECs (A)** Unbiased clustering of data from Uniform manifold approximation and projection (UMAP) of snATACseq (**B**) Transcriptional start site (TSS) enrichment score as defined by the ENCODE project (top) and fragment length (bottom) from the snATACseq (**C**) Top 6 enriched motifs in the LEC cluster. (**c’**) Probability of Tn5 insertion across the genome centered around predicted ETV2 binding motif in the LEC cluster (cluster 14) and in the macrophage cluster (cluster 19) have a higher enrichment compared to the mesothelial cells (cluster 7,23). (**D**) FeaturePlots illustrating high expression (gene activity) from the PROX1, NOTCH1, NR2F2 and SOX18 (Upper panels) and Motif enrichments for PROX1, RBPJ, NR2F2 and SOX18 (Lower panels). (**E**) Normalized probability of Tn5 insertion across the genome centered around predicted PROX1 and RBPJ binding motif (Top), and NR2F2 and SOX18 (Bottom). NR2F2 motifs show an increase in the TN5 insertion enrichment in the LECs compared to the other populations. (**F**) Top 6 enrichment motifs for the macrophage cluster. (**G**) Top 6 enrichment motifs for the mesothelial clusters.

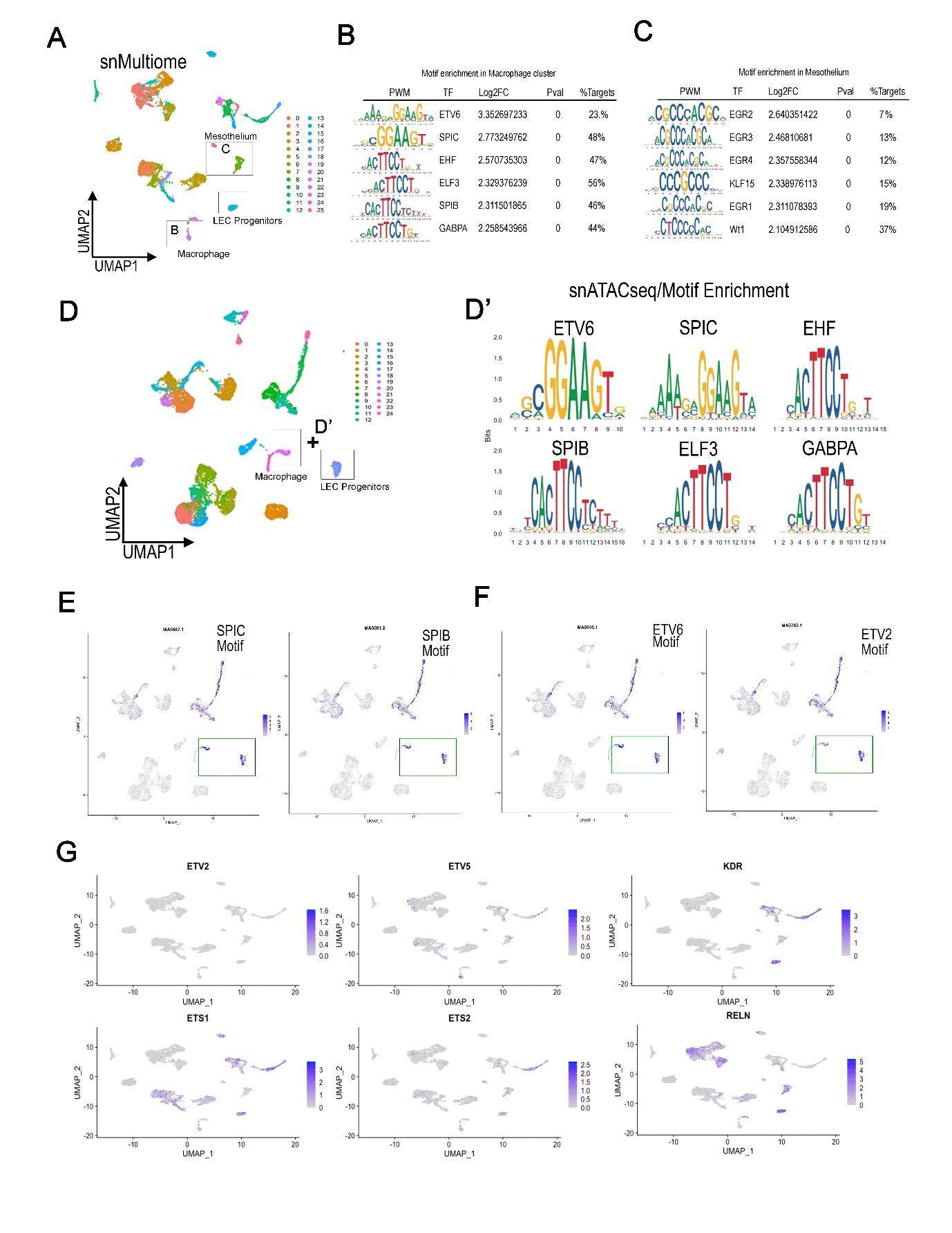
 **Suppl. Fig.4 Genomic comparison of macrophage and LEC clusters** (**A**) UMAP of single-Nuclei ATAC sequencing. (**B, C**) Top 6 enrichment motifs from the snMultiome macrophage cluster **(B)** and from the Mesothelium cluster (**C**). (**D**) UMAP of snATACseq illustrating the joint enrichment motifs analysis between the Macrophage and LEC clusters. (**D’**) The top 6 shared binding motifs identified ETV6 and other ETS transcription factors. (**E**) FeaturePlots depicting high Motif enrichments (signal) for the SPIC SPIB and ETV6 motifs in both clusters. In the same manner ETV2 motif enrichment shows a similar pattern. (**F**) FeaturePlots illustrating RNA expression from the WNN analysis of the snMultiome for ETS1, ETS2, ETV2, ETV5, KDR and RELN genes.
